# HIV transmission efficiency through contaminated injections in Roka, Cambodia

**DOI:** 10.1101/136135

**Authors:** David Gisselquist

## Abstract

A nosocomial HIV outbreak recognized in late 2014 in Roka commune, Cambodia, demonstrates the potential for rapid transmission through skin-piercing healthcare procedures. Information reported from the investigation of the Roka commune outbreak is sufficient to estimate the transmission efficiency of HIV through contaminated injection equipment. With conservative assumptions, two estimates are 4.6% and 9.2%. These estimates are much greater than widely disseminated and influential low estimates of risk from unsafe injections, estimates which have encouraged low estimates of the contribution of unsafe healthcare to Africa’s generalized HIV epidemics. More information about nosocomial risks in Roka commune could improve the estimates in this paper and advise HIV prevention programs, particularly in countries with unreliably sterile healthcare and high HIV prevalence.

## Introduction

In November 2014, a 74-year man and two relatives, all residents of Roka commune, Cambodia, tested HIV-positive.[1] Alerted to unexpected risks, other residents went for tests. Within weeks, more than 200 had tested HIV-positive.[2] In mid-December, government of Cambodia with assistance from the United States’ Centers for Disease Control and Prevention (CDC), the World Health Organization (WHO), UNAIDS, UNICEF, and Pasteur Institute initiated an investigation.[3,4] Through early 2017, investigators published selected findings in two reports.[5.6]

## Methods

Using information from two recent reports from the Roka commune outbreak, I calculate the estimated transmission efficiency of HIV through contaminated injection equipment during the Roka outbreak.

## Results

In Roka commune, Saphonn and colleagues collected and analyzed information on self-reported bloodborne and sexual risks in 112 HIV-infected cases and 214 HIV-negative controls. They report significant adjusted odds ratios (AORs) for infection associated with injections, infusions, and blood draws in the previous 0–6 months, infusions received 7 to 12 months previously, and injections received 13–36 months previously (Table 1). They also report positive but non-significant crude odds ratios (ORs) for HIV infection associated with obstetric procedures, major surgery, and minor surgery but not with tattooing or dental care during various periods in the previous 0–36 months.[6]

**Table 1:**
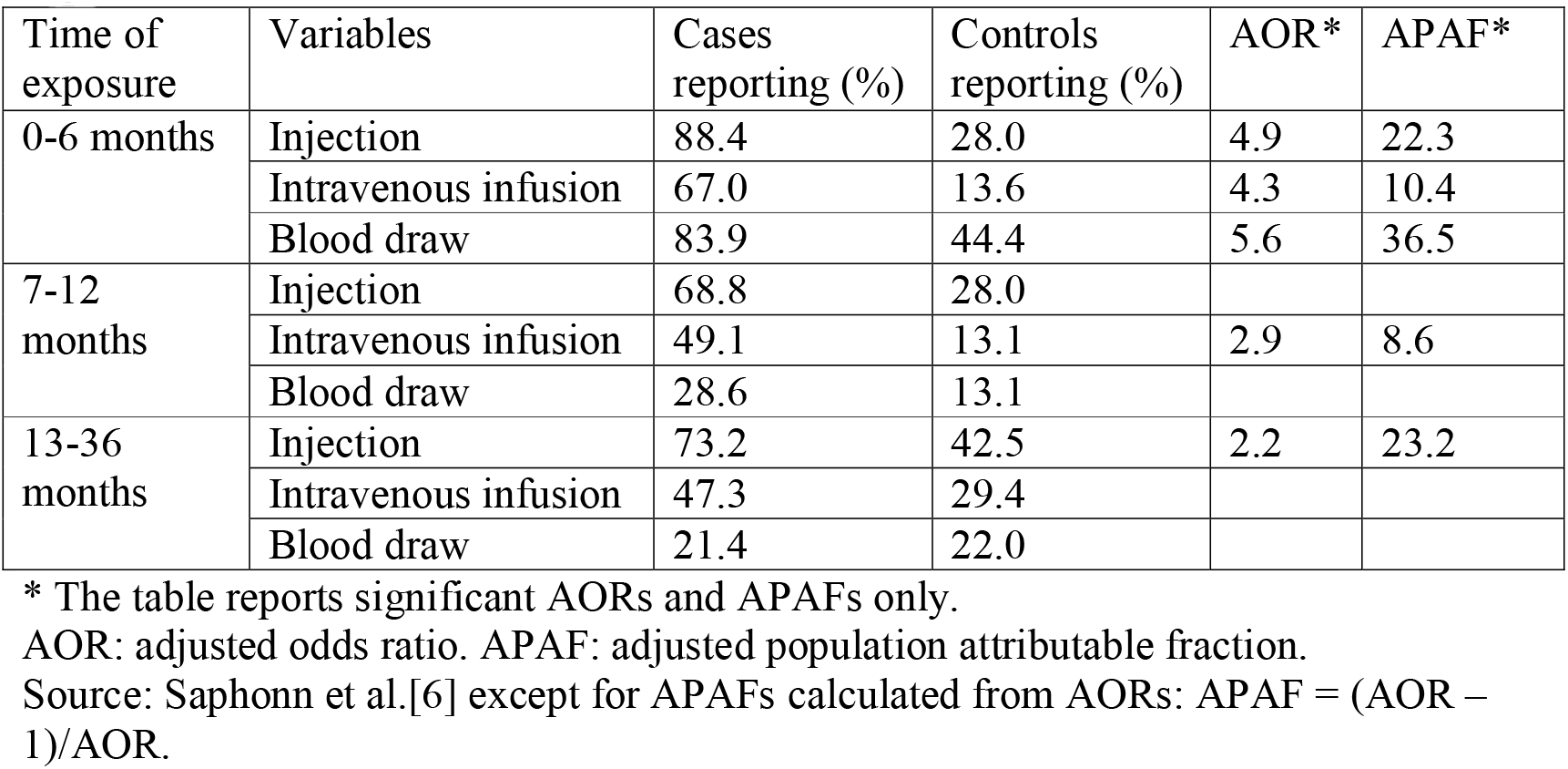
Population attributable fractions of HIV infections associated with blood exposures

| Time of exposure | Variables | Cases reporting (%) | Controls reporting (%) | AOR* | APAF* |
| --- | --- | --- | --- | --- | --- |
| 0-6 months | Injection | 88.4 | 28.0 | 4.9 | 22.3 |
|  | Intravenous infusion | 67.0 | 13.6 | 4.3 | 10.4 |
|  | Blood draw | 83.9 | 44.4 | 5.6 | 36.5 |
| 7-12 months | Injection | 68.8 | 28.0 |  |  |
|  | Intravenous infusion | 49.1 | 13.1 | 2.9 | 8.6 |
|  | Blood draw | 28.6 | 13.1 |  |  |
| 13-36 months | Injection | 73.2 | 42.5 | 2.2 | 23.2 |
|  | Intravenous infusion | 47.3 | 29.4 |  |  |
|  | Blood draw | 21.4 | 22.0 |  |  |

\* The table reports significant AORs and APAFs only.
AOR: adjusted odds ratio. APAF: adjusted population attributable fraction.
Source: Saphonn et al.[6] except for APAFs calculated from AORs: $APAF = (AOR - 1)/AOR$ .

Calculating from AORs, adjusted population attributable fractions (APAF) for injections, infusions, and blood draw in the previous 0–6 months are 22.3%, 10.4%, and 36.5%, respectively. Notably, the APAF for injections in the last 0–6 months is 32.2% (= 22.3/69.2) of summed APAFs for the three major risks during those months (69.2 = 22.3 + 10.4 + 36.5). The 36.5% APAF for blood draw 0–6 months before the study may have been inflated by people with recognized risks going for HIV tests and by people with HIV-positive tests from finger-stick blood going for confirmatory tests; notably, much higher percentages of cases than controls reported injections and infusions for all time periods, whereas for blood draw the difference falls to nil 13–36 months previously (Table 1). The authors report no OR or AOR greater than one for any sexual risk and no evidence of transmission from injecting illegal drugs, from mother-to-child, or from child-to-breastfeeder.

From this, I estimate injections account for at least a third – and likely closer to half – of recent infections, with infusions, blood draw, obstetric procedures, and major and minor surgeries accounting for most of the rest. To be conservative, subsequent calculations assume injections account for a third of recent infections.

Vun and colleagues[5] report, inter alia: (a) preliminary results from an incidence assay estimate 30% of infections acquired in the previous 130 days; (b) 242 residents of Roka commune tested HIV-positive during November 2014-February 2015; and (c) Cambodians with HIV infection receive an average of 7.2 injections per year (citing Cambodia’s 2005 Demographic and Health Survey).

From these data an estimated 73 (30% of 242) persons acquired HIV within 130 days before testing. To estimate transmission efficiency, I assume injections account for 24 (33% of 73) transmissions in the previous 130 days, that all blood draws were taken on the same day, that transmission rates were stable over the previous 130 days (i.e., that HIV prevalence increased exponentially from 169 to 242 persons for an average of 204 infected and potentially transmitting persons over the period), and that HIV-positive persons received an average of 2.56 injections in 130 days (at the rate of 7.2 per year). With these assumptions, all HIV-positive persons considered together received 522 injections (2.56 per person for the average 204 HIV-infected persons) in the previous 130 days.

With no information about sterilization practices, I consider two scenarios: that injection equipment was discarded or sterilized after 50% or 0% of injections administered to HIV-positive persons. If so, injection equipment was reused after 522 or 261 injections (100% or 50% of 522) administered to HIV-positive persons, infecting 24 subsequent persons, for a transmission efficiency of 4.6% (=24/522) of 9.2% (=24/261). These estimates consider the risk contaminated equipment from one injection administered to an HIV-positive person transmits HIV to one or more subsequent injectees, including persons receiving injecta from contaminated multi-dose vials.[7]

These estimates would be increased by considering that injections accounted for more than a third of infections, that some subsequent injectees were already infected, that some persons newly infected did not yet have HIV in their blood and so could not transmit, and that equipment was sterilized or discarded after more than 50% of injections administered to HIV-positive persons.

## Discussion

The estimates in this note have been developed with limited information about skin-piercing procedures in Roka commune. Unpublished details from the investigation might be able to improve these estimates. However, considering that 30% of infections were acquired in the previous 130 days, revised estimates with more detailed information are unlikely to be much less than what has been calculated in this paper and could be higher. Vun and colleagues[5] cite a previous estimate[7] of transmission efficiency through contaminated injections of 2%–7% from an outbreak in Romania, which roughly agrees with these estimates from Roka commune.

Over the last several decades, three teams assisted by WH0[8–10] have estimated transmission efficiencies of HIV through injections ranging from 0.3% to 1.2%, and have used these estimates to calculate healthcare injections accounting for 0.4%–4.2% of HIV incidence in Africa for various years from the late 1990s to 2010 (Table 2).

**Table 2:** Modeled estimates of the proportion of Africa’s HIV incidence from healthcare injections and estimated transmission efficiencies through injections used in the models

| Reference | Transmission efficiency through injections used to calculate incidence from injections (%) | Estimated HIV incidence from injections in (%) |  | Incidence estimated for which year |
| --- | --- | --- | --- | --- |
|  |  | Africa | World |  |
| Kane et al.[8] | 0.5%* | 1.25-2.5%† | 1.4-2.8%† | Late-1990s |
| Hauri et al.[9] | 1.2% | 2.5% | 5.0% | 2000 |
| Pepin et al.[10] | 0.32-0.64% | 2.1-4.2%‡ | 4.6-9.1% | 2000 |
|  | 0.32-0.64% | 0.4-0.8%‡ | 0.7-1.3% | 2010 |

\* Kane et al. estimate a transmission efficiency of HIV through injections of 0.3%, but their model uses 0.5% to estimate numbers of infections from injections.
† Kane et al. estimate 50,000-100,000 infections from injections in Africa and 80,000-160,000 in the world for an unspecified year in the late 1990s, equivalent to 1.25%-2.5% of UNAID's estimated 4 million incident infections in Africa in 1999[11] and 1.4%-2.8% of UNAIDS' estimated 5.8 million incident infections in the world in 1997.[12]
‡ Pepin and colleagues' estimate of 8,000-16,000 infections from injections in WHO's Africa regions AFR D and AFR E in 2010 and 53,000-106,000 in 2000. These estimates are 0.4%-0.8% of 2.2 million incident infections in Africa in 2009 and 2.1%-4.2% of 2.5 million incident infections in 2000 (derived from data in the UNAIDS' report Pepin cites).[13]

All estimates reported in Table 2 are based on evidence of transmission efficiency through needlestick accidents among healthcare workers, except that Pepin and colleagues’ consider as well two reports estimating HIV transmission efficiency among persons injecting illegal drugs (without noting the studies proposing those estimates assumed injection drug users shared equipment more often than reported).[14–16] None of the estimates of transmission efficiency reported in Table 2 consider evidence from investigated outbreaks.

Notably, with transmission efficiencies 0.3%–1.2%, an HIV-positive person would need an average of 83–333 injections to infect one other person (or more injections if some contaminated equipment was discarded or sterilized). With such low transmission efficiencies, neither the Roka commune outbreak or other recognized nosocomial outbreaks infecting more than 100 to thousands in Russia, Romania, China, Libya, Kazakhstan, Kyrgyzstan, and Uzbekistan[17] would have been possible.

Influential low estimates of HIV transmission efficiency through injections and related models estimating the percentage of HIV incidence in Africa from contaminated injections have deflected attention from nosocomial risk. Vun and colleagues note that “HIV transmission through medical injections has not historically been prioritized in Cambodia’s national HIV prevention strategy…”[p 143 in reference 5] A similar assessment applies to Africa, where silence about nosocomial risks has been common in HIV prevention messages and research. For example, a 2012 survey of high school students aged 12->20 years in rural South Africa reported more than 50% of infected boys (21 of 38) and girls (56 of 104) had never had sex; the authors considered under-reporting of sexual behavior but did not mention bloodborne risks.[18]

Information from the Roka commune outbreak could guide the general public and healthcare professionals to recognize and limit nosocomial risks. Such information could have the biggest impact on HIV prevention in countries with generalized epidemics and unreliably sterile healthcare. Several challenges are to provide details about unsafe practices suspected or identified during all the various skin-piercing procedures that likely contributed to the Roka commune outbreak and to improve estimates in this paper. Estimates might be improved by using more collected but unpublished data and by tracing and linking some of the infections.

## Acknowledgements

I thank Devon Brewer, John J. Potterat, John Sonnabend, Simon Collery, and Dan Gisselquist for thoughtful suggestions on analysis and presentation. I received no funding for this study and have no conflict of interest with respect to this article.

